# Autophagy protein Atg7 is essential and druggable for maintaining malaria parasite cellular homeostasis and organelle biogenesis

**DOI:** 10.1101/2023.08.16.553492

**Authors:** Akancha Mishra, Pratik Narain Srivastava, Shabeer Ali H, Satish Mishra

**Affiliations:** Division of Molecular Microbiology and Immunology, CSIR-Central Drug Research Institute, Lucknow 226031, India; Academy of Scientific and Innovative Research (AcSIR), Ghaziabad 201002, India

**Author notes:** Correspondence: Satish Mishra, Division of Molecular Microbiology and Immunology, CSIR-Central Drug Research Institute, Lucknow 226031, India. Faculty of Veterinary Medicine, University of Calgary, Calgary, T2N4N1, AB, CA.

**Keywords:** ATG7, ATG8, apicoplast, ER, autophagy, *Plasmodium*, malaria, sporozoites, liver stage, drug target

## Abstract

*Plasmodium* parasites have a complex life cycle that transitions between mosquito and mammalian hosts and undergoes continuous cellular remodeling to adapt to various drastic environments. Following hepatocyte invasion, the parasite discards superfluous organelles for intracellular replication, and the remnant organelles undergo extensive branching and mature into hepatic merozoites. Autophagy is a ubiquitous eukaryotic process that permits the recycling of intracellular components. Here, we show that the *P. berghei* autophagy-related E1-like enzyme Atg7 is expressed in the blood and liver stages, localized to the parasite cytosol and is essential for the localization of Atg8 on the membrane and the development of parasite blood and liver forms. We found that depleting Atg7 abolishes exocytosis of micronemes, organelle biogenesis and the formation of merozoites during liver stage development. Furthermore, we identified the compounds from the Maybridge library with a high docking score against *Pf*Atg7. We show that these compounds inhibit apicoplast biogenesis and parasite development in both blood and liver stages. Overall, this study establishes the essential functions of autophagy in *Plasmodium* blood and liver stages and highlights the potential of using Atg7 as a drug target against malaria.

## Introduction

The sporozoites invade hepatocytes and undergo extreme metamorphosis by completely dismantling the inner membrane complex (IMC), eliminating the invasive secretory organelles, and achieving a round trophozoite form ^1 2^. The early trophozoites undergo rapid mitotic division and divide their nucleus several times, and the remnant organelles undergo extensive branching and mature into hepatic merozoites. These merozoites are released in the bloodstream in the form of merosomes, where upon rupture, each merozoite invades and divides in RBCs ^3 4^. Throughout this development process, the parasite must eliminate unnecessary superfluous organelles to create room for newly synthesized stage-specific organelles and proteins. Understanding this process may provide insights into the parasite’s smooth transitions and development in different stages.

Autophagy (macroautophagy) is one such catabolic process widely conserved among eukaryotes and is induced in response to numerous cellular stresses, such as starvation, hypoxia, cellular differentiation, protein metabolism, invading pathogens, and aging ^5 6 7 8 9^. This involves the isolation of cytoplasmic components in a double membranous vesicle known as an autophagosome that later fuses and delivers its content to lysosomes for degradation. Lipids, amino acids, and other macromolecules generated after degradation in lysosomes are later recycled to maintain cellular homeostasis. Approximately 35 autophagy-related genes that have been characterized thus far in yeast are required for maintaining autophagy at various steps ^10^. Bioinformatic studies found partial conservation of ATG genes in apicomplexan parasites. Parasites lack the genes required for upstream sensing and signaling for the induction of autophagy but have genes needed for membrane expansion and completion. The redundant set of ATGs has also diversified their role more toward sustenance and aiding in extensive replication of the parasites in the host.

Atg8, a marker of active autophagy in yeast and mammals, is a small ubiquitin-like molecule, and its conjugation with phosphatidyl ethanolamine (PE) on the autophagosome membrane is essential for cargo selection, membrane tethering, hemifusion, expansion, and closure of autophagosomes ^11 12 13 14^. In apicomplexan parasites, Atg8 is conserved along with the intermediate genes required for its conjugation with PE and is specifically localized on the apicoplast membrane. The discrete localization of Atg8 on the apicoplast membrane throughout the parasite life cycle must have a distinct role from its role in autophagy. In a recent study, the loss of Atg8 function led to the complete loss of apicoplast and arrested parasite growth in *P. falciparum* ^15^. Similar defects in apicoplast biogenesis were observed in *T. gondii* Atg8 knockdown, which led to a block in parasite replication ^16^. Targeting Atg7 (E1 activating enzyme), which is required for Atg8 conjugation, resulted in attenuation of parasite growth in *P. falciparum* blood stages ^17^. Similar phenotypes of growth attenuation were observed in *TgAtg4* and *TgAtg3* knockdown, depicting the essentiality of the pathway in parasite survival.

Recent reports suggest a role for Atg8 in *P. falciparum* blood stage development; however, the importance of the Atg8 conjugation system in liver stage development remains largely unknown ^17^. In this study, we demonstrate the role of the autophagy-related protein Atg7 in the *P. berghei* life cycle with a special focus on its role in liver stage development. The Atg7 gene was found to be indispensable in *P. berghei* blood stages. By utilizing Flp/FRT-based conditional mutagenesis, we elegantly demonstrate its role in Atg8 conjugation, clearance of micronemes, organelle biogenesis, and expansion of parasites during liver stage development. We identified Atg7-specific inhibitors and showed that compounds inhibit *Plasmodium* blood and liver stage development by inhibiting parasite autophagy machinery.

## Results

### Atg7 is predominantly expressed during asexual blood and liver stages and localized to the parasite cytosol

We began our study by evaluating the spatiotemporal localization of Atg7 at various stages of parasite development. For this, we generated a transgenic parasite line that endogenously expresses the *Atg7*-3XHA-mCherry fusion protein (Fig. S1A and S1B). The *Atg7*-3XHA-mCherry parasite line completed the life cycle normally, indicating that Atg7 was unaffected by tagging (Fig. S1C and S1D). For expression analysis, blood-stage parasites were immunostained with anti-mCherry and anti-Hsp70 antibodies. mCherry signals showed the cytosolic localization of Atg7 in the schizont, ring, and trophozoite stages, with PCC values of 0.672, 0.929, and 0.929, respectively (Fig. 1A). For liver stage expression analysis, HepG2 cells were infected with *Atg7*-3XHA-mCherry sporozoites, and EEFs harvested at different time points were stained with anti-mCherry and Hsp70 antibodies. IFA revealed cytosolic localization with PCC values of 0.775 and 0.444 at 24 and 40 hpi, respectively (Fig. 1B). The mCherry signals were not detected 55 hpi, revealing the early to mid-liver stage expression of Atg7 protein. mCherry signals were not observed during mosquito stages (data not shown).

**Figure 1.**
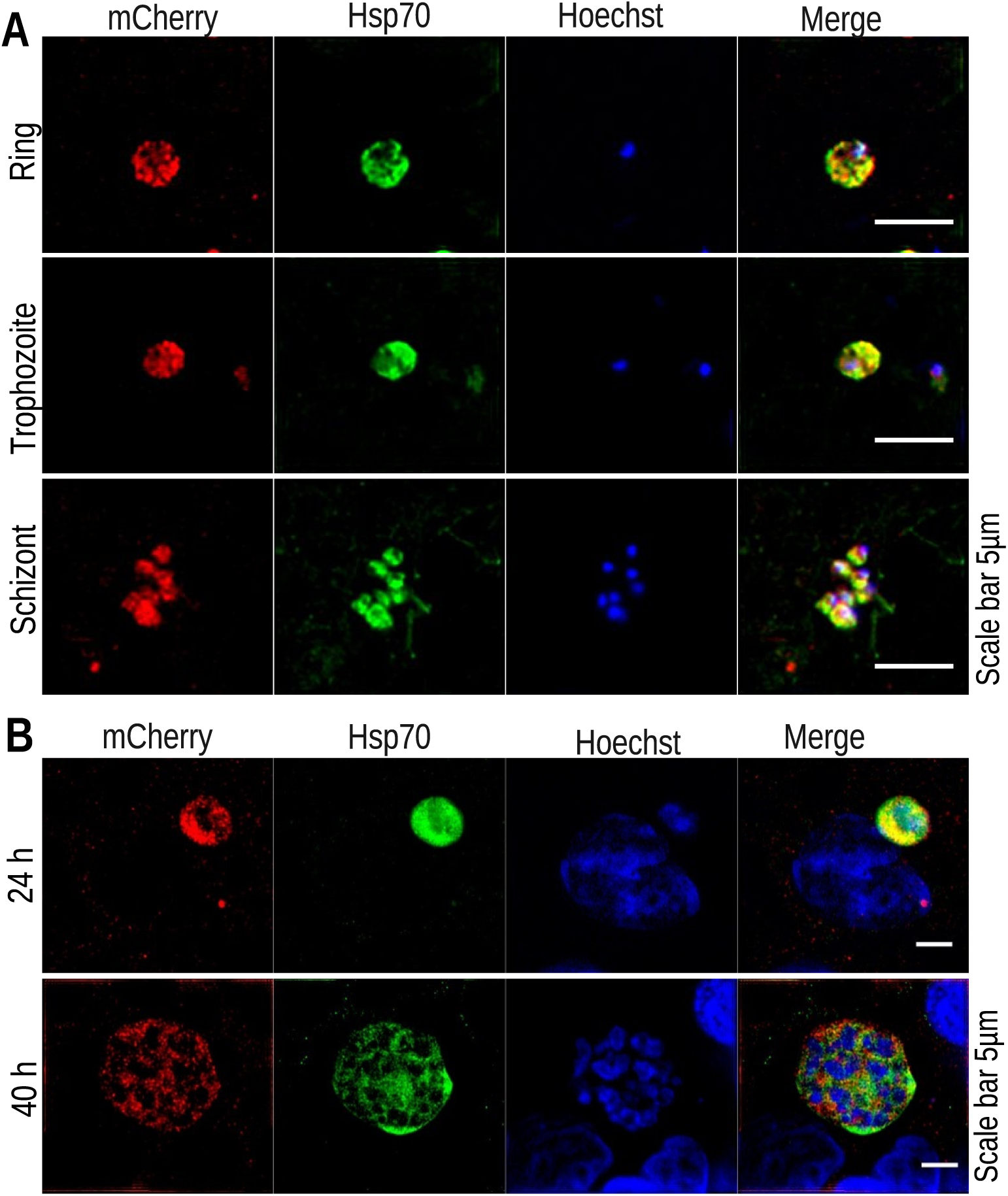
Atg7 expression and localization. **(A)** Atg7-3XHA-mCherry-infected blood stage slides were prepared, fixed, and immunostained with anti-Hsp70 and mCherry antibodies. Merged images showed the colocalization of Atg7 with HSP70 in the ring, trophozoite and schizonts, indicating the cytosolic expression and localization of Atg7 in blood stages with PCC values of 0.672, 0.929 and 0.929, respectively. Scale bar 5 µm. **(B)** HepG2 cells infected with Atg7-3XHA-mCherry sporozoites were fixed at 24 and 40 hpi and stained with anti-Hsp70 and mCherry antibodies. Similar to blood stages, we found that Atg7 was expressed in the EEF cytosol. The Atg7 and Hsp70 signals colocalized at both time points (24 and 40 hpi) with PCC values of 0.775 and 0.444, respectively. The nuclei were stained with Hoechst 33342.

### Atg7 is essential for the development of *P. berghei* blood and liver stages

To decipher Atg7 function in the *Plasmodium* life cycle, we attempted to directly disrupt Atg7 three times by double crossover homologous recombination but failed to obtain KO parasites in the blood stage (Fig. S2A). Next, we employed the yeast Flp/FRT-based conditional mutagenesis system to conditionally silence Atg7 function in sporozoites ^18^. We engineered two FRT sites in such a way that after excision in mosquito stages, a portion of the gene was excised (Fig. S2B-S2E). To ensure that the excision of the flirted *Atg7* locus was specific to the TRAP/FlpL line, the construct was also transfected into *P. berghei* WT parasites (Fig. S2F). These genetic manipulations did not affect parasite development in the blood and mosquito stages (Fig. S3A-S3E). To check the excision efficiency of the *Atg7* gene in sporozoites, genotyping was performed, which revealed successful excision of the flirted *Atg7* locus transfected in the TRAP/FlpL line but not in the *P. berghei* WT parasites, which do not express FlpL (Fig. S3F). These results suggest that Atg7 is not required for the progression of sporozoites from the midgut to the salivary gland. To analyze the role of Atg7 in liver stage development, sporozoites were intravenously injected into C57BL/6 mice, and the prepatent period was observed. We found that all the mice inoculated with TRAP/FlpL or *Atg7* cKO/WT sporozoites became patent on day 3 p.i., whereas parasites lacking Atg7 (*Atg7* cKO) failed to initiate blood-stage infection (Fig. 2A and Table 1). We observed rare blood stage infection, and genotyping revealed a nonexcised *Atg7* locus (Fig. S3F). This indicates that few parasites that escape the excision of their flirted locus in the mosquito stages were able to complete liver stage development and initiate blood stage infection. To identify the stage-specific roles of Atg7, C57BL/6 mice were injected i.v. with sporozoites, and parasite burden in the liver was quantified by real-time PCR analysis of *Pb*18S rRNA transcripts. We found comparable parasite burden in *Atg7* cKO and TRAP/FlpL parasites until 38 hpi, which was decreased in late time point samples harvested at 55 hpi (Fig. 2B). As the parasites undergo schizogony, they start expressing merozoite surface protein 1 (MSP1), whose product is important for merozoite formation. We found a significant decrease in MSP1 transcripts in *Atg7* cKO parasites at 55 hpi (Fig. 2C). During exoerythrocytic schizogony, parasites undergo multiple rounds of nuclear division, increase in size, and mature into infectious merozoites that fill the sac called the merosome. To visualize this development pattern in *Atg7* cKO parasites, HepG2 cultures were infected with sporozoites and harvested at different time points. We found that *Atg7* cKO sporozoites invaded hepatocytes normally (Fig. S3G and S3H). The parasites in the infected HepG2 cells were quantified using qRT‒ PCR, which was consistent with the in vivo liver infectivity (Fig. 2D). Immunofluorescence analysis of exo-erythrocytic forms (EEFs) with Hsp70 and UIS4 antibodies followed by quantification revealed normal development until 24 hpi. However, EEFs harvested at 40, 55, and 64 hpi showed arrested growth and a reduction in size and number (Fig. 2E-G). On quantifying the Hoechst-stained DNA centers, we also observed a strong defect in nuclear fragmentation in Atg7 cKO parasites at late time points and a significantly increased nuclear/EEF area fraction (Fig. 2H-L).

**Figure 2.**
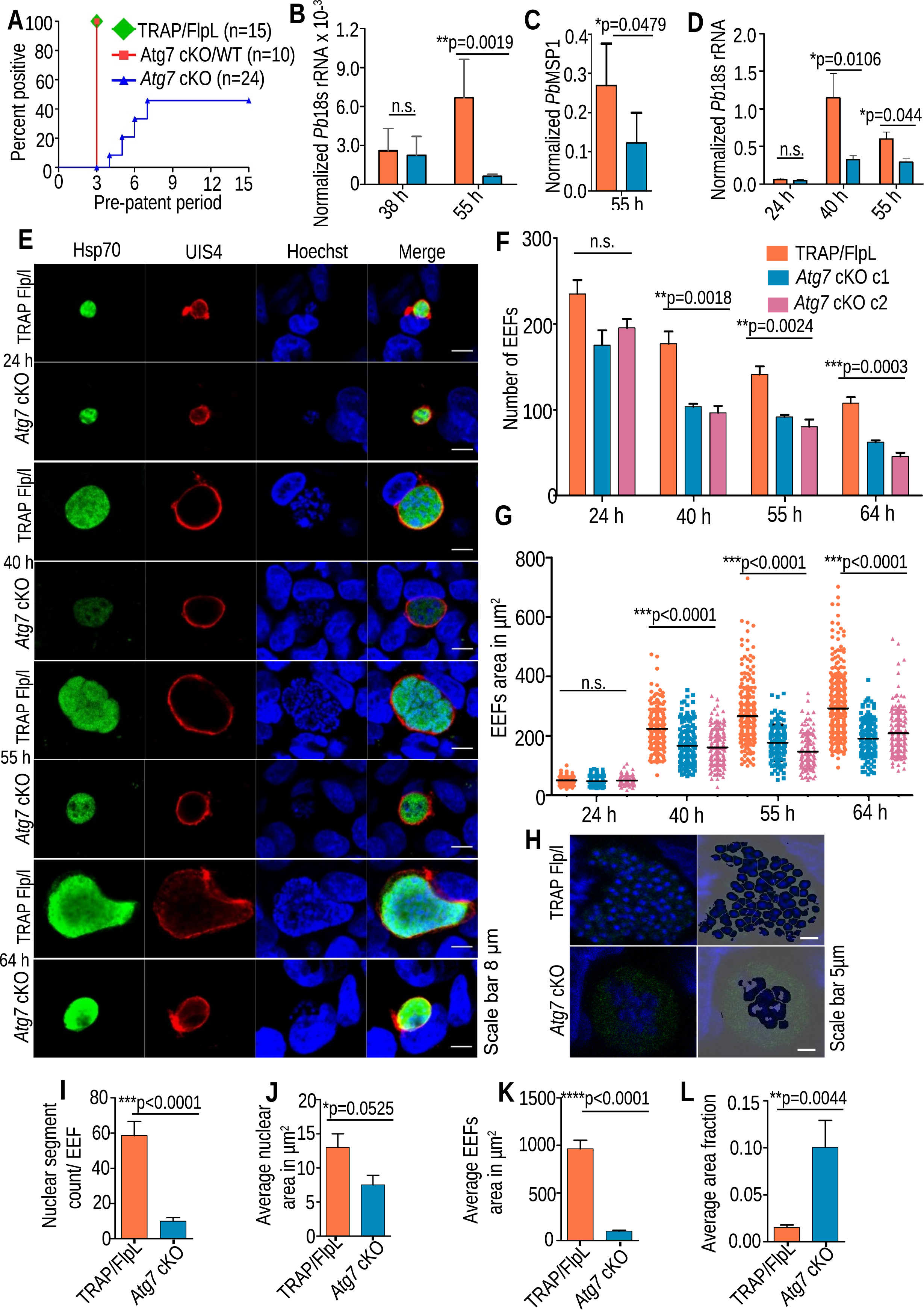
Atg7 is essential for liver stage development. **(A)** Infectivity of *Atg7* cKO sporozoites in C57BL/6 mice. All mice injected with *Atg7* cKO/WT or TRAP/FlpL parasites became patent on day 3. The mice that became patent in the Atg7 cKO group carried a nonexcised *Atg7* locus, as confirmed by genotyping. **(B)** Real-time analysis of parasite burden in the liver showing the comparison of normalized *Pb*18s rRNA transcript numbers in TRAP/FlpL- and *Atg7* cKO-infected mice at 38 and 55 hpi. Representative of three independent experiments is presented as the mean ± SD, n=5 (no significant difference, 38 h (p=0.9933), significantly different 55 h (p=0.0019), Student’s t test. **(C)** Quantification of *PbMSP1* transcript numbers in TRAP Flp/L- and *Atg7* cKO-infected mouse livers harvested at 55 hpi. Representative of three independent experiments is presented as the mean ± SD, n=5 (statistically significant (p=0.0479), Student’s t test. **(D)** Quantification of parasite burden in infected HepG2 cells harvested at 24, 40, and 55 hpi. Data are presented as the mean ± SEM, n=3 biological replicates (no significant difference at 24 h (p=0.6194), significant difference at 40 h (p=0.0106), and 55 h (p=0.044), Student’s t test). **(E)** HepG2 cells infected with *Atg7* cKO or TRAP Flp/L sporozoites were fixed at 24, 40, 55, and 64 h.p.i. and analyzed by IFA. Parasites were stained with anti-UIS4 and anti-Hsp70 antibodies. Nuclei were stained with Hoechst 33342. **(F)** The number of EEFs was quantified and is presented as the mean ± SEM. n=3 biological replicates (no significant difference at 24 h (p=0.0760), significant difference at 40 h (p=0.0018), 55 h (p=0.0024), and 64 h (p=0.0003), one-way ANOVA). **(G)** Measurement of EEF area at 24 h, 40, 55, and 64 hpi. Data were pooled from three independent experiments. n=150 to 200 EEFs per parasite strain (no significant difference at 24 h p=0.3055. significant difference at 40 h, 55 h, and 64 h (p<0.0001), one-way ANOVA**. (H)** *Atg7* cKO parasites have compromised nuclear segmentation. TRAP/FlpL and *Atg7* cKO EEFs fixed at 64 h were identified by the Hsp70 antibody, and the nucleus was stained with Hoechst 33342. **(I)** Bar graph showing decreased nuclear segments, **(J)** decreased average nuclear **(K)** and EEF area **(L)**, and increased average nuclear/EEF area fraction in *Atg7* cKO parasites in comparison to TRAP/FlpL EEFs. Data are presented as the mean ± SEM. n=16, from 3 biological replicates. (Significant difference (p<0.000026), (p<0.0525), (p<0.000022), (p<0.0044), Student’s t test).

**Table 1.** Infectivity of *Atg7* cKO sporozoites in C57BL/6 mice. Mice were i.v inoculated with *Atg7* cKO, TRAP/FlpL or Atg7 cKO/WT sporozoites.

| Experiment | Parasites | Number of sporozoites injected | Mice positive/Mice injected | Pre-patent period (Days) |
| --- | --- | --- | --- | --- |
| 1 | TRAP/FlpL | 5000 | 3/3 | 3 |
|  | *Atg7* cKO SC1 | 5000 | 4/5 | 5.75 |
| 2 | TRAP/FlpL | 5000 | 3/3 | 3 |
|  | *Atg7* cKO SC1 | 5000 | 1/5 | 5 |
| 3 | TRAP/FlpL | 5000 | 3/3 | 3 |
|  | *Atg7* cKO SC1 | 5000 | 1/5 | 6 |
| 4 | TRAP/FlpL | 5000 | 3/3 | 3 |
|  | *Atg7* cKO/WT | 5000 | 5/5 | 3 |
|  | *Atg7* cKO SC2 | 5000 | 4/5 | 5.5 |
| 5 | TRAP/FlpL | 5000 | 3/3 | 3 |
|  | *Atg7* cKO/WT | 5000 | 5/5 | 3 |
|  | *Atg7* cKO SC2 | 5000 | 1/4 | 5 |

### *Atg7* cKO parasites failed to mature into hepatic merozoites due to impaired organelle biogenesis

Arrested growth of the *Atg7* cKO EEFs was further confirmed by observing merozoite development by staining with MSP1 antibody, which was found to be impaired, including the nuclei count (Fig. 3A and B). After the maturation of parasites in the liver, they are released in the form of merosomes ^4^. We enumerated the merosome numbers in TRAP/FlpL; however, due to the absence of merosomes in the *Atg7* cKO parasites, the collected supernatant was divided equally and injected into five Swiss mice (Fig. 3C). The TRAP/FlpL group injected with 10 merosomes became patent on day 5; however, all the mice injected with *Atg7* cKO culture supernatant remained negative during the observation period (Fig. 3D and Table S1). These findings indicated that in the absence of Atg7, the parasite’s DNA replication and segregation were impaired, and the parasites failed to develop into infective merozoites, which explains their inability to initiate blood stage infection. Atg8 localization on endomembranes confers many unique roles in their mammalian counterparts, such as membrane tethering, hemifusion, and expansion of autophagosome membranes ^12^. However, both autophagosomes and apicoplasts originate in the endomembrane system, and the disruption of Atg8 in *P. falciparum*/*T. gondii* results in a defect in apicoplast biogenesis ^15 19 20 21 22^. To assess whether loss of Atg7 function had a role in organelle biogenesis during liver stage development, apicoplasts and ER were visualized using IFA. We found extensive branching of the apicoplast and ER in TRAP/FlpL parasites, while this branching was lost in the *Atg7* cKO parasites, suggesting impaired development of organelles in the mutant parasites (Fig. 3E and F).

**Figure 3.**
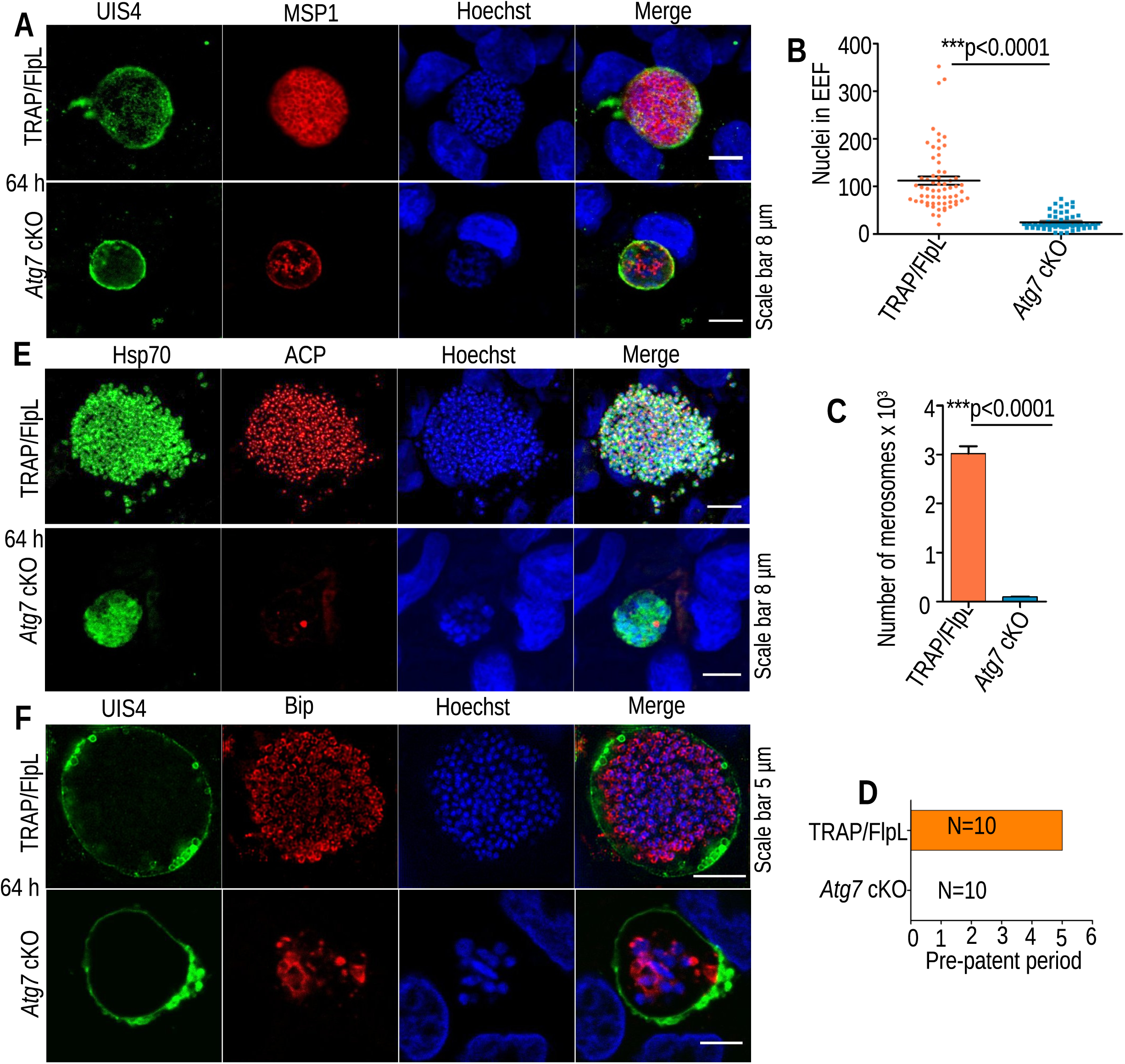
*Atg7* cKO parasites show impaired organelle biogenesis and fail to mature into hepatic merozoites. **(A)** *Atg7* cKO or TRAP/FlpL EEFs fixed at 64 hpi were stained with MSP1 and UIS4 antibodies. TRAP/FlpL mature EEFs showed clear nuclear segregation, with each nucleus surrounded by MSP1, whereas the *Atg7* cKO parasites failed to segregate their nuclei, and MSP1 development was impaired. **(B)** Nuclei count in *Atg7* cKO or TRAP/FlpL EEFs. Hoechst-stained nuclei images were acquired under a confocal microscope, and nuclei were counted using an ImageJ cell counting tool. Data were pooled from three independent experiments (n=70). (Significant difference, (P<0.0001), Student’s t test). The nuclei were stained with Hoechst 33342. (**C)** The number of detached cells (merosomes) in *Atg7* cKO or TRAP/FlpL-infected HepG2 cultures. Data are presented as the mean ± SEM. n=3 biological replicates. (Significant difference (p<0.0001), Student’s t test). (**D)** Merosome formation was impaired in *Atg7 c*KO parasites. Swiss mice were injected intravenously with merosomes or culture supernatant that failed to initiate blood stage infection. **(E and F)** EEFs fixed at 64 hpi were immunostained with ACP/Hsp70 and Bip/UIS4 antibodies. EEFs showed impaired organelle biogenesis, and both apicoplast and ER development were aborted in *Atg7* cKO parasites.

### Atg7-mediated conjugation of Atg8 is required for biogenesis of the apicoplast

With significant growth defects observed at the late stages of exo-erythrocytic schizogony, we next analyzed the Atg8 distribution pattern in the late liver stages of parasitic development. *Atg7* cKO and TRAP/FlpL EEFs fixed at 64 hpi were immunostained with anti-ACP and anti-Atg8 antibodies to visualize Atg8 lipidation on the branched apicoplast structures. We found Atg8-mediated apicoplast expansion and division in TRAP/FlpL parasites, and this conjugation and branching were completely lost in *Atg7* cKO parasites (Fig. 4A and B). These results indicate that due to the lack of Atg8 conjugation, the apicoplast failed to expand and segregate during schizogony.

**Figure 4.**
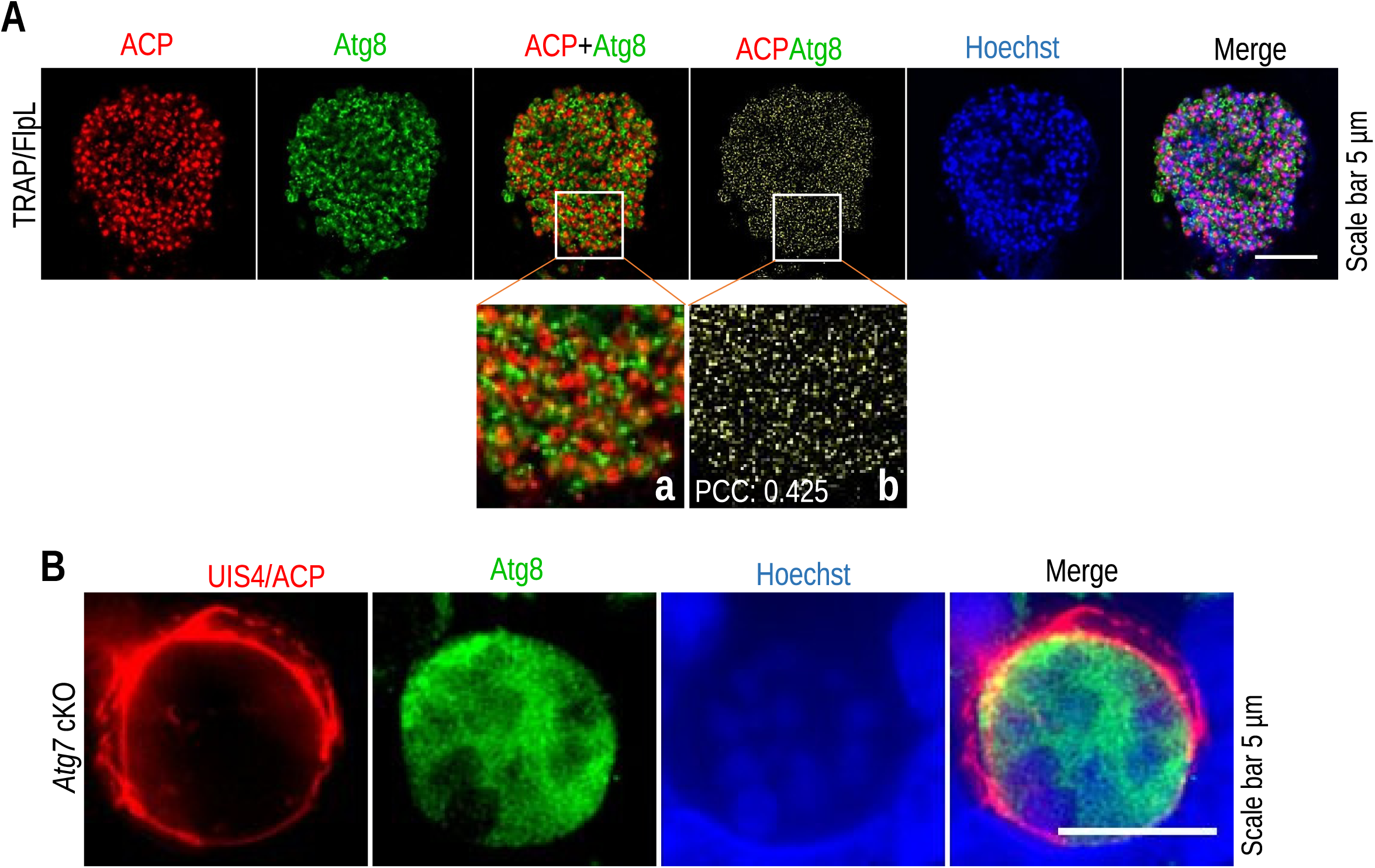
Atg8 localization on the membrane is required for apicoplast branching. (**G)** Atg7 cKO parasites failed to conjugate Atg8 to the apicoplast. TRAP/FlpL EEFs harvested at 64 hpi were immunostained with anti-ACP and anti-Atg8 antibodies. A significant portion of Atg8 was found to be localized on the apicoplast (PCC: 0.425). Inset images **(a, b)** show a zoomed portion of conjugated Atg8 on the apicoplast. The nuclei were stained with Hoechst 33342. **(H)** *Atg7* cKO EEFs harvested at 64 hpi were immunostained with anti-ACP and anti-Atg8 antibodies. EEFs were identified using an anti-UIS4 antibody. The KO parasites failed to conjugate Atg8 and expand the apicoplast. The nuclei were stained with Hoechst 33342.

### Atg7 is essential for the exocytosis of unnecessary superfluous organelles during liver stage development

During parasite development in the liver stage, a series of morphological changes occur that involve the mechanical elimination of the secretory organelles, which are no longer required posthepatocyte invasion by the parasite. Previous studies have reported the presence of membrane whorls enclosing the IMC, rhoptries, and micronemal contents secreted out in the extracellular matrix, which are further degraded by the proteolytic enzymes released by the parasite ^23^. Here, we monitored the elimination pattern of micronemes by using TRAP antibodies during the liver stage development of *Atg7* cKO and TRAP/FlpL parasites. We first started with its identification in sporozoites and then followed its track in EEFs. We found the same distribution pattern of micronemes in TRAP/FlpL and *Atg7* cKO parasites at 0 h (Fig. 5A). By 6 to 12 hpi in TRAP/FlpL parasites, micronemal content was found to accumulate in the center of EEFs as the parasite achieved spherical morphology (Fig. 5A). *Atg7* cKO parasites also showed a similar distribution pattern, but there was a significant increase in the intensity of TRAP signals at 12 hpi as calculated by CTCF (Fig. 5B). TRAP/FlpL EEFs observed 24 hpi showed TRAP-enriched organelles migrating toward the PV membrane, while they remained in the center in the *Atg7* cKO parasites. (Fig. 5A). TRAP/FlpL EEFs fixed at 40 and 55 hpi showed 70 to 83% elimination of micronemal content, while it was approx. 36 and 40% in *Atg7* cKO parasites. Quantification of TRAP signals revealed an increase in CTCF values post-12 h, indicating a severe defect in the exocytosis of these rudimentary organelles in *Atg7* cKO parasites (Fig. 5B).

**Figure 5.**
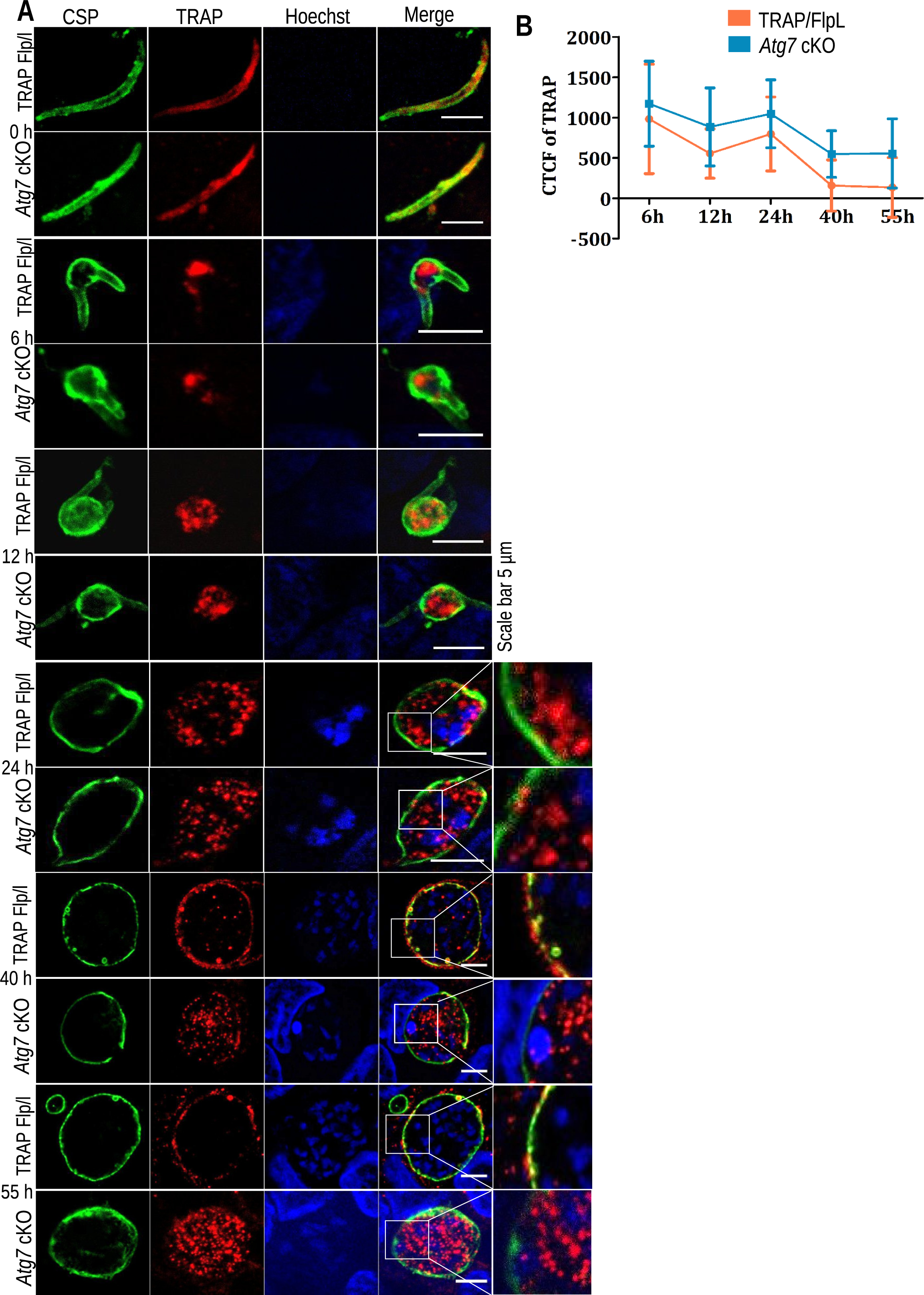
*Atg7* cKO parasites failed to exocytose micronemes during liver stage development. **(A)** HepG2 cultures infected with *Atg7* cKO or TRAP/FlpL sporozoites were fixed at different time points. The EEFs were stained with TRAP antibody to track the movement of micronemes and stained with CSP antibody to identify the EEFs. The images showed the aggregation of the micronemes by 6 to 12 hpi, and by 24 h, the micronemes began to divert toward the PV membrane of the parasites in TRAP/FlpL EEFs, while in *Atg7* cKO parasites, the micronemes remained centered. By 40 to 55 hpi, most of the micronemal content was excreted out of the parasite cytosol, while a significant number of *Atg7* cKO EEFs retained their micronemes. **(B)** Quantification of TRAP fluorescence signals in TRAP/FlpL or *Atg7* cKO EEFs fixed at 6-55 hpi. The EEFs were stained with TRAP antibody, and its signal inside the PV membrane was calculated as corrected total cell fluorescence (CTCF) (n= 30-35 EEFs). The nuclei were stained with Hoechst 33342.

### Atg7 as a drug target and its inhibitor kills *P. falciparum* blood stages

The promising phenotype of *Atg7* knockout parasites in both the blood and liver stages has led us to explore it as a drug target. Atg7 is an E1-activating enzyme that adenylates Atg8 in an ATP-dependent manner. The *Pf*Atg7 protein has a similarity of 34% with *Hs*Atg7 and 32% with *Sc*Atg7. To begin with inhibitor design, the ATP binding site was targeted, and the initial redocking experiment indicated that AutoDock Vina could successfully reproduce the crystal structure pose of ATP in its binding site in 3VH4 (*Sc*Atg7)^24^. The same docking parameters were employed to dock ATP to the *Pf*Atg7 and *Hs*Atg7 models (Fig. 6A). Similar poses were produced in both cases, indicating the validity of the docking protocol. Residues involved in ATP binding were analyzed to decipher the binding site for further virtual screening. LYS944, ASP967, ILE968, ASP1003, SER1007, CYS1177, and THR1178 in *Pf*Atg7 and LYS413, MET435, SER436, ILE437, SER480, ASP476 and CYS572 in *Hs*Atg7 were found to be possibly involved in ATP binding (Fig. S4). Differences in the type of binding residues indicate a possibility of finding a selective ligand that can inhibit *Pf*Atg7 while not binding to *Hs*Atg7, hence being antimalarial while not being toxic to the host. The Maybridge library was screened for compounds with high docking scores with *Pf*Atg7. A docking score cutoff of 7.5 was chosen to filter the initial list of docked compounds, as this was the docking score of ATP in the *Pf*Atg7 binding site. The top 100 hits with docking scores higher than 7.5 were selected from this step and screened for binding with HsATG7 using the same protocol and parameters. Using the same criteria as above, but in reverse, the top 15 compounds were selected that had docking scores lower than 7.5 with HsATG7 (Table S2). The compounds were screened for antimalarial activity against the *P. falciparum* asexual blood stages, and compounds RJF01701 and BTB01219 were found to be active. Both compounds bind *Pf*Atg7 differently, and major portions of both molecules occupy the Atg7 tunnel-shaped cavity adjacent to the ATP binding site. The cavity is surrounded by LEU1001, ASP1003, LYS1005, AGR1008, VAL1134, ALA1135, ILE1136, TYR1156, CYS1177 and SER1185, providing more improved binding support than ATP (Fig. 6B and C). Both compounds showed different modes of binding with *Hs*Atg7 (Fig. 6D and E). The tunnel adjacent to the ATP binding site remains unoccupied, which makes the interaction less stable despite the possibility of one hydrogen bond formation, which is not found in the case of *Pf*Atg7 (Fig. 6D and E). We found that compounds RJF01701 and BTB01219 inhibited *P. falciparum* development with IC50 values of 2.26 and 5 µM, respectively (Fig. 6F, G and S5). Both compounds were further screened for toxicity in HepG2 and Vero cell lines and found to be not toxic at or approximately 70 µM. These results demonstrate the activity of both compounds against *Pf*Atg7 while being nontoxic to humans. To elucidate that the compounds inhibit *Plasmodium* autophagy protein Atg7, we attempted to express recombinant *Pf*Atg7 to setup a drug assay but failed. Alternatively, we observed the conjugation pattern of Atg8 and apicoplast development. We found that treated parasites were mostly arrested during the trophozoite stage and failed to undergo schizogony. Atg8 was also found to be mostly in the unconjugated form in the cytosol, and apicoplast branching was impaired (Fig. 6H and Fig. S6). The results indicated that the compounds possibly inhibited *Pf*Atg7, leading to the failure of Atg8 conjugation to the apicoplast membrane, which affected parasite development.

**Figure 6.**
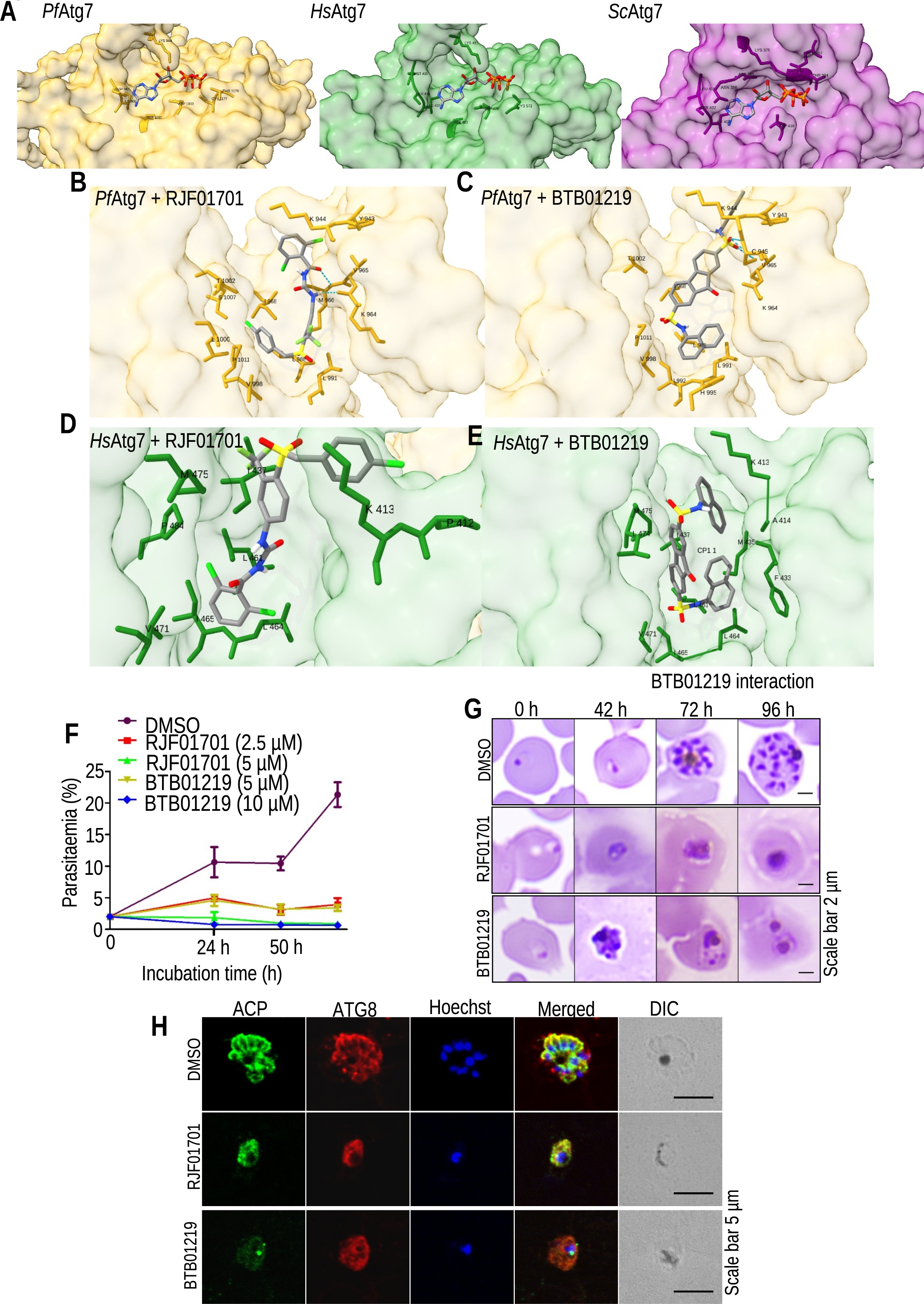
Atg7 structure model generation, screening of the compound library and parasite development inhibition assay. **(A)** Residues involved in ATP binding in *Pf*ATG7, HsATG7, and ScAtg7 (3VH4). *Pf*ATG7 – yellow, HsATG7 – green, *Sc*Atg7 – magenta. **(B)** Giemsa-stained blood smear showing aborted parasite development in the RJF01701- and BTB01219-treated groups harvested at 0, 42, 72, and 96 h posttreatment. **(B)** Different binding poses of compound RJF01701 (2,6-dichloro-N-[[4-[(4-chlorophenyl)methylsulfonyl]-3-(trifluoromethyl)phenyl]carbamoyl]benzamide). **(C)** and compound BTB01219 (2-N,7-N-dinaphthalen-1-yl-9-oxofluorene-2,7-disulfonamide) in the ATP binding site of *Pf*Atg7. The amino acids of the domain involved in the binding are highlighted in yellow, forming stable hydrogen bonds with the compounds highlighted in gray. **(D)** Binding poses of compound RJF01701 **(E)** and compound BTB01219 in the ATP binding site of *Hs*Atg7. No stable hydrogen bonds are formed between residues of the targeted site highlighted in green and the compounds highlighted in gray. **(F and G)** Time course of *P. falciparum* 3D7 asexual blood stage parasite growth inhibition by compounds RJF01701 and BTB01219 in comparison to the DMSO control. Data are presented as the mean ± SEM, n=3 biological replicates (significant growth difference between DMSO and RJF001701 and BTB01219 treated at IC50, (p=0.0210), one-way ANOVA), significant growth difference between DMSO and RJF001701 and BTB01219 treated at 2X of IC50, (p=0.0062), one-way ANOVA). **(H)** IFA with ACP and Atg8 antibodies of *P. falciparum* asexual blood stage parasites treated with DMSO, RJF01701, and BTB01219 showed impaired apicoplast branching and Atg8 conjugation in treated parasites. The nuclei were stained with Hoechst 33342.

### Characterization of *P. berghei* liver stage inhibition using Atg7 inhibitors

Next, we evaluated the activity of both compounds against the *P. berghei* liver stages using an in vitro luciferase assay. The compound RJF01701 inhibited liver stage development with an IC50 of 7.53 µM at 50 hpi but not at 24 hpi (Fig. 7A). The area of EEFs was comparable in control and treated samples harvested at 24 hpi, whereas a significant attenuation in EEF size was observed at 50 hpi (Fig. 7B). Surprisingly, compound BTB01219 did not show a significant decrease in luciferase activity (Fig. 7C); however, the size of EEFs was found to be reduced at 50 hpi (Fig. 7B). Due to the differential activity of both compounds against the two parasite strains, the compounds were docked using the same docking parameters as used previously against *Pf*Atg7, and we found that RJF01701 and BTB01219 interact differently at the ATP binding site of *Pb*Atg7 (Fig. 7D and E). The compound RJF01701 formed a more stabilized bond at the cavity by forming a hydrogen bond with Met26 (Fig. 7D and F); however, no such H-bonds were formed with BTB01219 (Fig. 7E and F). Both compounds were also very closely apposed with *Pb*Atg7 in comparison to *Pf*Atg7 or *Hs*Atg7, which suggested its even stronger binding with *Pb*Atg7 (Fig. 7G). The anti-liver stage activity of RJF01701 mimics the phenotype of *Atg7* cKO parasites, suggesting that the identified compounds inhibit the autophagy pathway of malaria parasites, leading to late liver stage attenuation.

**Figure 7.**
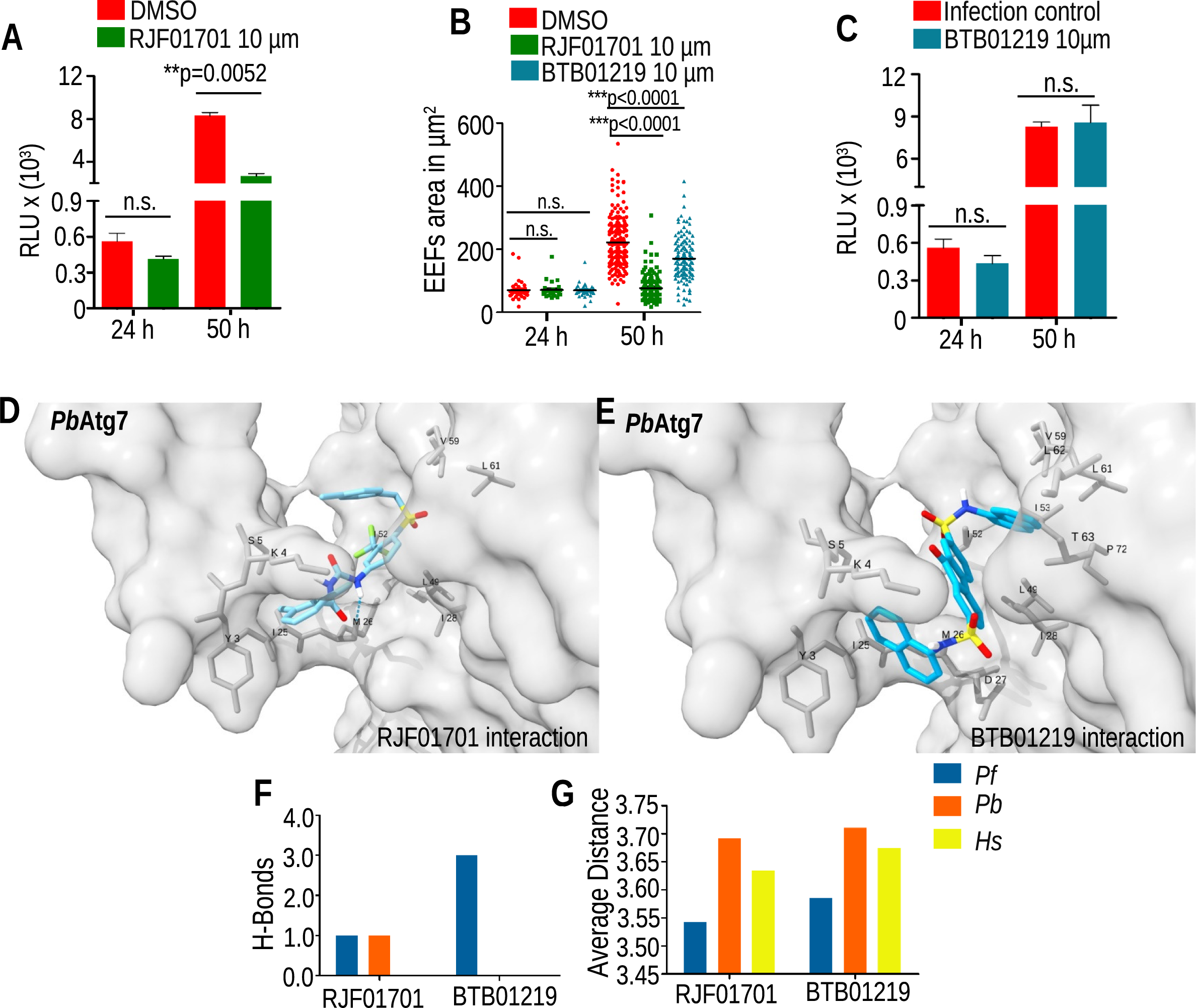
Compound RJF01701 inhibits *P. berghei* liver stage development. **(A)** Luciferase activity of *P. berghei* Luc-infected HepG2 cells. Bioluminescence was estimated by the addition of D-luciferin at 24 and 50 hpi. Data representative of two independent experiments are presented as the mean ± SD (significant difference in RLU between DMSO control and parasites treated with RJF01701 harvested at 50 hpi (p=0.0052, Student’s t test), whereas no significant inhibition was observed when samples were harvested at 24 hpi (p=0.1089, Student’s t test). **(B)** Comparison of the EEF area between treated and DMSO controls harvested at 24 and 50 hpi. Data represent values from two independent experiments (significant difference between control and parasites treated with RJF01701 and BTB01219 at 10 µM, (p<0.0001), Student’s t test). **(C)** Luciferase activity of *P. berghei* Luc-infected HepG2 cells. Bioluminescence was estimated by the addition of D-luciferin at 24 and 50 hpi. Data representative of two independent experiments are presented as the mean ± SD. No significant inhibition was observed between the DMSO control and parasites treated with BTB01219 (24 hpi (p=0.2042) and 50 hpi, (p=0.8428), Student’s t test). **(D)** ATP binding pose of *Pb*Atg7 shows the difference between the residues involved in the binding of RJF01701 **(E)** and BTB01219. The interaction of RJF01701 with *Pb*Atg7 was stabilized by one hydrogen bond, while no such stable bonds were formed with BTB01219. This indicates its better binding and in vitro activity. **(F)** Comparison between hydrogen bonding of *Pf*Atg7, *Pb*Atg7, and *Hs*ATG7 with compounds. RJF01701 has one hydrogen bond each with P*f*Atg7 and *Pb*Atg7, while BTB01219 has one hydrogen bond with *Pf*Atg7. However, neither has a hydrogen bond with *Hs*ATG7. **(G)** Comparison between the average distance between the compounds and the target site. Both compounds have a lower average contact distance with *Pf*Atg7 compared to both *Pb*Atg7 and *Hs*ATG7. A lower distance indicates stronger binding and may indicate better in vitro activity.

## Discussion

In this study, we show that *Pb*Atg7, which is an intermediate enzyme of the Atg8 conjugation pathway, plays an essential role in parasite blood and liver stages. *Atg7* cKO parasites failed to exocytose micronemes, undergo organelle biogenesis, and mature into hepatic schizonts. Compounds identified through in silico docking inhibited *Plasmodium* blood and liver stages. Our finding is consistent with previous studies showing the essentiality of the Atg8 conjugation pathway in *P. falciparum* asexual blood stages ^17 25 15^. Many autophagy studies on apicomplexans have shown the discrete localization of Atg8 on the apicoplast, and loss of Atg8 directly affects its biogenesis ^19 20 21 26 22 16^.

Apicoplast localization of Atg8 in apicomplexan parasites has diversified its role apart from autophagy. We have demonstrated the direct involvement of Atg8 in the biogenesis of apicoplasts in late liver stages. *Atg7* cKO parasites that failed to conjugate Atg8 on apicoplasts resulted in the complete loss of apicoplast expansion and division and failed to mature into hepatic merozoites that did not initiate blood stage infection in mice. This implies that Atg8 localization on endomembranes might be a source of membrane supply for the extensive branching of apicoplasts during the replicative phase of parasite development. This was also supported by a study in which overexpression of Atg8 led to abnormally fast expansion and branching of the apicoplast ^27^. Another study on *T. gondii* showed that apicoplast genome segregation is synchronized with nuclear division and that the apicoplast interacts with centrosomes during division ^28^. This interaction might be mediated via Atg8, as its downregulation led to improper segregation of apicoplasts into individual tachyzoites ^16^. Although *Plasmodium* parasites lack centrioles, they have a centrosome-like structure, and Atg8 might interact with it, which enables the positioning of the apicoplast during division ^29 30^.

The existence of apicoplasts has been proven to be crucial for parasite survival. The organelle has lost its photosynthetic abilities but has retained several important metabolic pathways, such as the biosynthesis of fatty acids, isoprenoids, heme, and iron-sulfur clusters ^31^. Loss of the integrity of the apicoplast or interference with any of its biosynthetic pathways would affect the supply of such important metabolites that affect parasite development. In a study, the loss of apicoplast protein PALM resulted in parasite attenuation in the liver stage that further failed to develop into merozoites and did not initiate blood stage infection ^32^. Similar detrimental effects on growth were observed when genes of the FASII pathway were deleted, leading to parasite arrest in liver stage forms ^33 34 35^. Impaired apicoplast expansion hampers the subcellular distribution and supply of lipids that might indirectly affect ER differentiation, as suggested by their closed apposition and vesicle sharing ^36^.

Furthermore, our study implicates the role of Atg7 in the maintenance of cellular homeostasis via a special secretory autophagy pathway present in the *Plasmodium* parasite. In this pathway, organelle-enclosed endomembranous vesicles are exocytosed by a secretory pathway and later degraded by an extracellular enzymatic process ^37 38 39^. We found that TRAP-filled micronemal contents tend to move toward the PV membrane and be exocytosed by 40-55 hpi in wild-type parasites; however, this process was impaired in *Atg7* cKO parasites, suggesting the role of the Atg8 conjugation pathway in the elimination of secretory organelles during liver stage development ^1 27^. Mammalian homologs of Atg8, LC3, and GABARAP have been shown to interact with microtubules through their LIR motifs, which might enable the movement of autophagosomes and GABA receptor-like vesicles ^40 41 42^. Similarly, Atg8 on micronemal vesicles might interact with microtubules and enable the movement and elimination of the vesicle out of the parasite lumen.

Overall, in our study, we showed the importance of the Atg8 conjugation pathway in *P. berghei* blood and liver stage development. The pathway enables the clearance of micronemal vesicles from the parasite cytosol and has also been diversified and plays a role in the biogenesis of organelles during exoerythrocytic schizogony. Finally, our identified compounds against *Plasmodium* Atg7 inhibit blood and liver stages and hold the potential to develop a multistage drug to alleviate malaria.

## Materials and Methods

### Parasites and cell lines

*Plasmodium berghei* ANKA (MRA 311) was obtained from BEI resources, USA. The marker-free transgenic *P. berghei* ANKA line expressing FlpL under *Plasmodium* stage-specific promotor-TRAP was used for the generation of conditional knockout parasites^18^. *Plasmodium* sporozoites were obtained by infecting female *Anopheles stephensi* mosquitoes as described previously^18^. *A* chloroquine-sensitive *Plasmodium falciparum* strain (3D7, MRA-151) was used to evaluate the antimalarial activity of the compounds (BEI Resources, USA). Parasites were maintained in fresh human erythrocytes at 2% hematocrit in complete RPMI-1640 (Sigma) medium in addition to 0.5% AlbuMaxII (Gibco by Life Technologies), 0.2% glucose (Merck Life Science), 0.2% NaHCO_3_ (SD Fine Chemical Limited) and 15 µM hypoxanthine (Sigma) and incubated at 37°C in a 5% CO_2_ incubator ^43^. The parasites were regularly synchronized with 5% D-sorbitol^44^. Human liver hepatocellular carcinoma (HepG2) and Vero (African Green Monkey Kidney) cells (ATCC) were regularly maintained in DMEM (Sigma) supplemented with 10% FBS (Sigma), 0.2% NaHCO_3_ (Sigma), 1% sodium pyruvate (Genetix), and 1% penicillin‒streptomycin (Invitrogen) at 37°C with 5% CO2. We routinely tested cell lines for mycoplasma contamination.

### Mice

Female Swiss albino mice (6-8 weeks old) were used for routine parasite infections and passages, and C57BL/6 mice 6-8 weeks old were used for sporozoite in vivo infectivity. All animal procedures conducted were approved by the Institutional Animal Ethics Committee at CSIR-Central Drug Research Institute, India (approval no: IAEC/2013/83 and IAEC/2018/3).

### Generation of transgenic and knockout parasites

For the generation of *Pb*Atg7-3XHA-mCherry transgenic parasites, two fragments, F1 (0.66 kb) and F2 (0.62 kb), were amplified using primers 1632/1633 and 1634/1324 and sequentially cloned and inserted into the *Pb*C-3XHA-mCherry hDHFR plasmid at *Xho*I*/Bgl*II and *Not*I*/Asc*I (NEB), respectively. The plasmid was linearized with *Xho*I*/Asc*I and transfected into *P. berghei* ANKA purified schizonts as described previously ^45^. In an attempt to generate direct knockout of *Pb*Atg7, two fragments, F3 (0.54 kb) and F4 (0.61 kb), were amplified using primers 1321/1322 and 1323/1324 and sequentially cloned and inserted into pBC-GFP-yFCU-hDHFR at blunt-ended *Sal*I and *Not*I/*Asc*I *sites,* respectively. The plasmid was linearized with *Xho*I/*Asc*I restriction digestion and transfected into *P. berghei* ANKA schizonts^45^. For the generation of conditional knockout of *Pb*Atg7, three fragments, F5 (1.03 kb), F6 (0.16 kb) and F7 (0.62 kb), were amplified using primers 1325/1326, 1327/1328 and 1353/1330, respectively. All three fragments F5, F6, and F7 were sequentially cloned and inserted into a p3’TRAP-flitre-hDHFR plasmid at *Hind*III*/Not*I, EcoRV, and *Sal*I*/Kpn*I restriction sites, respectively. For swapping the 3’UTR of the gene with TRAP, 12 bp was added to the F6 fragment for the continuation of 3’UTR function. The cloned plasmid was linearized by restriction digestion with *Hind*III*/Kpn*I and transfected into *P. berghei* ANKA TRAP/FlpL purified schizonts ^18 46^. All transfected parasites were selected by oral administration of pyrimethamine (0.07 mg/ml, Sigma) for 6 consecutive days, and resistant parasites were collected and genotyped. For 5’ and 3’ integrations, the primers 1399/1216 and 1215/1400 were used for *Atg7* cKO and primers 2022/1218 and 1215/1400 for Atg7-3XHA-mCherry transgenic parasites, respectively. All primers used for cloning and diagnostic PCRs are given in Table S3. Clonal lines of the cKO and transgenics were obtained by limiting dilutions of the parasites in Swiss mice and used for further analysis.

### Analysis of asexual blood stage propagation

To analyze the effect of swapping the 3’UTR of the gene with the TRAP 3’UTR during blood stage development, an equal number of iRBCs of Atg7 cKO and TRAP/FlpL parasites were i.v. injected into Swiss mice. The growth was monitored daily by making Giemsa-stained blood smears.

### Analysis of parasite development in the mosquito

For parasite transmission in the mosquito, female *Anopheles stephensi* mosquitoes were fed on Swiss mice infected either with *Atg7* cKO or TRAP/FlpL parasites and kept in an environmental chamber maintained at 19°C with 80% relative humidity. On day 14 postbloodmeal, the mosquito midgut was dissected and imaged under a Nikon Eclipse 80i microscope using a 10x (numerical aperture [NA], 0.25; air) objective with no filter, and the sporulation pattern and oocyst numbers were determined. To determine the oocyst sporozoite number, mosquito midguts were crushed and spun at 50 xg for 4 minutes, the supernatant was collected, and the sporozoite number was counted using a hemocytometer under a Nikon phase contrast microscope using a 40x (NA 0.75; air) objective with no filter. To achieve the optimal excision efficiency of the flirted *Atg7* locus by the flippase enzyme, mosquito cages were transferred to 25°C on day 17 post blood meal. On days 21-23, salivary glands were dissected, and the sporozoite number per mosquito was enumerated. To check the excised flirted locus, genomic DNA was isolated from sporozoites using a genomic DNA purification kit (Promega) and genotyped using primers 1409/1400 (primers are listed in Table S3) ^18^.

### Sporozoite in vivo infectivity

To assess the in vivo infectivity of sporozoites, C57BL/6 mice were i.v. injected with 5,000 sporozoites (5 mice/group). The onset of blood-stage infection was estimated by daily examination of Giemsa-stained blood smears. The patent mice were genotyped using primers 1409/1400. To estimate the liver stage parasite biomass, another group of C57BL/6 mice was i.v. inoculated with 5,000 *Atg7* cKO or TRAP/FlpL sporozoites, and the liver was harvested and homogenized at different time points in TRIzol reagent (Himedia). Total RNA was isolated using the manufacturer’s instructions. Randomly primed cDNA was synthesized using 1 µg of RNA in a reverse transcriptase reaction containing 1X PCR buffer, 0.5 mM dNTPs, 5 mM MgCl2, 20 U RNase inhibitor, 2.5 µM random hexamers and 50 U reverse transcriptase (Applied Biosystems) in a thermocycler (Eppendorf). The resulting cDNA was used for the quantification of *Pb*18s rRNA, *Pb*MSP1, and mouse GAPDH transcripts as described previously ^47^ using SYBR green reagent (Takara) with primer pairs 1195/1196, 1219/1220, and 1193/1194, respectively. The transcript number of parasites was expressed as the normalized value by taking the ratio of *Pb*18s rRNA/mGAPDH or *Pb*MSP1/mGAPDH. All the data were obtained using a CFX Opus 96 real-time PCR system (Bio-Rad), and the primers are listed in Table S3.

### Sporozoite in vitro infectivity

The hepatocyte infectivity and development assay was performed as previously described ^48^. Briefly, HepG2 cells were plated on 48-well or 24-well plate cultures at a density of 55,000 or 100,000 cells per well, respectively. For the invasion assay, salivary gland sporozoites (10,000 sporozoites/well) were added to a 48-well culture plate and allowed to invade for 2 hours. The cells were fixed with 4% paraformaldehyde (PFA) for 20 minutes at RT. For the development assay, salivary gland sporozoites (5,000 sporozoites/well) were added to the 48-well plate culture and maintained as described previously ^49^. The culture was fixed at 24, 40, 55, and 64 hpi using 4% PFA for 20 minutes at RT. For the merosome assay, salivary gland sporozoites (30,000 sporozoites/well) were added to the 24-well plate culture, and at 64 hpi, the culture was observed live. The supernatant was collected and quantified using a hemocytometer. To check merosome infectivity, Swiss mice were i.v. injected with harvested merosomes (10 merosomes/mouse). The onset of blood-stage infection was observed by Giemsa-stained blood smears. The genotyping of infected mice was performed using primers 1409/1400 (primers are listed in Table S3). To estimate the parasite biomass, HepG2 cultures infected in a 24-well plate were harvested in TRIzol reagent at different time points, RNA was isolated and reverse transcribed, and transcripts were quantified using real-time PCR as described above.

### Immunofluorescence assay

For the invasion assay, fixed cultures were washed twice in PBS and blocked with 1% BSA/PBS, and the extracellular and intracellular sporozoites were immunostained with anti-CSP mouse antibody ^50^ (diluted 1 µg/ml) before and after permeabilization as described previously^48^. Primary antibodies were then detected using anti-mouse IgG conjugated to Alexa Fluor 594 or Alexa Fluor 488 (diluted 1:500; Invitrogen), and nuclei were stained with Hoechst 33342 (Invitrogen), mounted using prolong diamond antifade reagent (Life Technologies) and visualized under a Nikon Eclipse 80i fluorescence microscope (100X (NA 1.30; oil) objective). For EEF development, fixed cultures were washed twice in PBS, permeabilized with 0.1% Triton X-100 (Sigma) for 10 minutes at RT, washed again in PBS and blocked with 1% BSA/*PB*S. Primary antibodies were diluted in 1% BSA/PBS and incubated for 1 h at RT, and then the wells were washed three times with PBS. Primary antibodies were then detected using anti-mouse or anti-rabbit Alexa Fluor-conjugated IgG as described above. To analyze the EEF number and area, EEFs were stained with Upregulated in infectious sporozoites gene 4 (UIS4) ^51^ (diluted 1:1,000, rabbit polyclonal) and heat shock protein 70 (Hsp70) ^52^ (diluted 1:500, mouse monoclonal). The EEFs were counted using a Nikon Eclipse 80i fluorescence microscope, and the area of the EEFs was measured using NIS-D software. Merozoite formation was visualized using merozoite surface protein 1 (MSP1) antibody ^53^ (diluted 1:500, mouse monoclonal). Apicoplast and ER development was visualized using acyl carrier protein (ACP) antibody (diluted 1:1,000, rabbit polyclonal ^54^) and Bip (diluted 1:500, rat polyclonal, BEI Resources), respectively. To visualize micronemes, a thrombospondin-related anonymous protein (TRAP) antibody was used. Affinity-purified polyclonal rabbit antibody against *P. berghei* TRAP was developed by GenScript Inc., Piscataway, NJ, against the peptide sequence CAEPAKPAEPAEPAE. For microneme distribution, EEFs were fixed at different time points and immunolabeled with TRAP (diluted 1:200, rabbit polyclonal) and CSP ^50^ (diluted 1:1,000, mouse monoclonal) antibodies. The TRAP and CSP signals were revealed using Alexa Fluor 594-conjugated anti-rabbit IgG and Alexa Fluor 488-conjugated anti-mouse IgG, respectively (diluted 1:500; Invitrogen). Nuclei were stained with Hoechst 33342, and the coverslips were mounted using Prolong Diamond antifade reagent (Life Technologies). Representative images were acquired using FV1000 software on a confocal laser scanning microscope (Olympus BX61WI) using a UPlanSAPO 100x (NA 1.4, oil) or 63x (NA 0.25, oil) objective or a Leica-SP8 confocal microscope (Leica, Wetzlar, Germany) under a 100X oil immersion (NA 1.4, oil) objective or a Leica DM 3000 LED microscope using a 100X (NA 1.25, oil) objective. Images were processed using ImageJ Fiji software and deconvoluted using the “Deconvolution Lab2” plugin.

### Immunolocalization

For localization studies, different parasite stages were fixed with 4% PFA (Sigma) for 20 min at RT. The parasites were permeabilized with chilled methanol at 4°C for 15 min, blocked with 1% BSA/*PB*S, and immunolabeled with anti-mCherry (diluted 1:500, Novus Biologicals) and anti-Hsp70 (diluted 1:500) antibodies for 2 h at RT. The mCherry and Hsp70 signals were revealed using Alexa Fluor 594-conjugated anti-rabbit IgG and Alexa-Fluor 488-conjugated anti-mouse IgG, respectively (diluted 1:500; Invitrogen). Nuclei were stained with Hoechst 33342. The images were taken using a Leica 3000 LED microscope and pseudocolored using ImageJ fiji “Lookup Table” features. The colocalization of the protein was estimated using the Pearson correlation coefficient calculated using the ImageJ “JACop” plugin.

### In silico 3D modeling and screening of a library

Three-dimensional structural models of *Pf*ATG7 and its closest human homolog, HsATG7 (UniProt ID: O95352), were accessed from the alpha-fold database. A full-length initial model of HsATG7 was used to avoid any potential bias in docking parameters. A crystal structure of the c-terminal domain of HsATG7 bound to ATG8 and ATP as a cofactor (PDB ID: 3VH4) was used as a template to identify potential ATP binding sites in both models by superposition. AutoDock Vina was used to redock the ATP molecule to 3VH4 to optimize docking parameters for generating a binding conformation similar to the ligand conformation in the crystal structure^24^. The docking parameters thus obtained were used for further high-throughput virtual screening (HTVS) and binding optimization experiments. ATP was also redocked to the *Pf*ATG7 and HsATG7 models to further optimize the respective binding sites. A docking search grid was created around these binding sites to reduce the search space and improve the speed of HTVS. All docking parameters were incorporated in a python script to automate the HTVS experiments. The Maybridge compound library (approximately 50000 compounds) obtained from Thermo Fisher was used to perform the HTVS experiments. Compound information was converted into 3D structures, completed by hydrogen addition and energy-optimized using the open babel toolkit. All compounds were rapidly docked on the *Pf*ATG7 and HsATG7 ATP binding sites. Top hits were estimated by assigning a cutoff based on the docking score of ATP. Compounds with high *Pf*ATG7 scores and low HsATG7 scores were selected for in vitro testing. This filtering strategy was employed to identify compounds that could be antimalarials by *Pf*ATG7 inhibition while at the same time not being toxic to the host by inhibiting HsATG7.

### In vitro assessment of anti-*Plasmodium* activity of compounds

The antimalarial activity of in silico-identified compounds against *Plasmodium falciparum* strain 3D7 blood stages was quantified using a SYBR Green I fluorometric assay ^55^. Briefly, the Maybridge library compounds (Thermo Fisher, USA) were twofold serially diluted in duplicate in 96-well plates (ranging from 5 µM to 312 nM concentrations), followed by the seeding of synchronized *Pf*3D7 ring stage culture at 1% parasitemia and 2% hematocrit in standard RPMI 1640 (Sigma)-based medium. Chloroquine (CQ) (Sigma) was used as a positive control in the range of 50 - 3.12 nM. The culture was maintained at 37°C in an incubator with 5% CO_2_. The culture was terminated by adding 100 µl lysis buffer with 20 mM Tris (Promega), 5 mM EDTA (Amreso), 0.008% saponin (Sigma), and 0.08% Triton X-100 (Sigma), and 2X SYBR Green (Thermo Fisher Scientific) dye was added to each well and incubated in the dark at RT for 30-45 min. Fluorescence signals were detected and quantified using a plate reader (Synergy HT, BioTek) at excitation 485 nm and emission 535 nm wavelengths, and the IC_50_ values were calculated. CQ was used as a standard, and infected and uninfected erythrocytes were used as positive and negative controls, respectively. The cultures obtained from the active compounds RJF01701 and BTB01219 were harvested at 40, 72, and 96 h post-treatment, and parasite growth was monitored by making Giemsa-stained blood smears. Blood smears were also immunostained with anti-Atg8 and anti-ACP antibodies to visualize apicoplast development and Atg8 distribution in the treated parasites. Images were acquired using FV1000 software on a confocal laser scanning microscope (Olympus BX61WI) using a UPlanSAPO 100x (NA 1.4, oil) objective and processed through ImageJ fiji.

### Cytotoxicity of compounds against mammalian cells

The cytotoxicity of the two lead compounds was tested in the Vero and HepG2 cell lines. Briefly, 1x10^4^ cells/well were seeded in a 96-well plate in medium containing DMEM, 10% FBS and 1% penicillin‒streptomycin and grown for 24 hr at 37°C in an incubator with 5% CO_2_. Doxorubicin (Sigma) was used as the standard drug. Untreated cells were used as the positive control, and only medium was used as the negative control. Cultures were incubated at 37°C for 72 hr in a 5% CO_2_ incubator. Next, 10 µl of MTT (Invitrogen) solution (5 mg/ml) was added to each well, and the plates were incubated at 37°C for an additional 2 h. The media containing MTT was removed from all the wells, and the crystals were dissolved by adding 100 µl DMSO (Sigma) per well. The fluorescence intensity was detected and quantified using a microplate reader (Epoch-BioTek) at excitation and emission wavelengths of 590 and 650 nm, respectively.

### *P. berghei* luciferase liver stage assay

For the luciferase assay, a parasite line expressing GFP-luciferase was generated by transfecting a plasmid into *P. berghei* as previously described^56^. Compounds with potential activity against *P. falciparum* blood stages were tested against *P. berghei* Luc-infected HepG2 cells. Briefly, HepG2 cells (30,000/well) were seeded in 96-well plates (Greiner Bio) and infected with 4,000 *P. berghei* Luc sporozoites. The culture was maintained as described above along with continuous treatment with 2-fold serially diluted (ranging from 20 µM to 2.5 µM) RJF0101 and BTB01901 compounds. Images of the GFP-expressing parasites were acquired using an EVOS cell imaging system (Invitrogen) at 24 and 50 hpi to evaluate the EEFs. To quantify the luciferase activity, D-Luciferin (Steady-Glo Luciferase Assay System, Promega) was added to each well, luciferase activity was detected using a GloMax Navigator (Promega), and the IC50 was determined. Uninfected cells incubated with D-luciferin and infected cells without D-luciferin were used as the negative control. Infected (DMSO) control wells were used as a positive control.

### Antibody generation

The *P. falciparum* Atg8 (PF3D7_1019900)-expressing clone was used for the expression of recombinant protein^57^. The pET32a-*Pf*Atg8 plasmid, which expresses Atg8 as a thioredoxin-His fusion (Trx-His-Atg8), was transformed into *E. coli* BL21 (DE3) cells, and expression was induced using isopropyl b-D-thiogalactopyranoside (SRL) and further purified as previously described ^57^. The rat was primed with purified protein in complete Freund’s adjuvant and boosted twice with incomplete Freund’s adjuvant. The rats were bled, and the serum was collected for further analysis.

### Nuclei count

The quantification of nuclei per EEF was performed using ImageJ Fiji software. TRAP/FlpL and *Atg7* cKO 64 hpi EEFs stained with UIS4 and Hoechst 33342 were used for quantification. Hsp70 staining was used to visualize the parasite, and the DNA centers were counted using the “multipoint” feature. For segmentation, the images were deconvoluted followed by analysis using the thresholding and watershed segmentation features of ImageJ.

### Statistical analysis

Data are presented as the means ± sems or mean ± sd. Statistical analysis was performed using GraphPad Prism Software. Statistical significance between the two groups was analyzed using an unpaired two-tailed t test or one-way analysis of variance (ANOVA). A P value less than 0.05 was considered statistically significant.

### Data and Materials Availability

All data are available within this manuscript, and raw data are available from the corresponding author upon reasonable request. Materials generated in this study are available from the corresponding author on request.

## Acknowledgments

We are thankful to Dr. Robert Menard (Institute Pasteur, France) for the p3’trap-hDHFR-flirte3 vectors. We thank Dr. Kota Arun Kumar (University of Hyderabad, India) for the modified version of p3’trap-hDHFR-flirte3, pBC-3XHA-mCherry-hDHFR vectors, and WT mCherry parasites. We thank Drs. Photini Sinnis (Johns Hopkins University, USA) and Leiden University Medical Center (The Netherlands) for the P. berghei TRAP/FlpL parasite line. We thank Dr. Anthony A. Holder (The Francis Crick Institute, UK), Drs. Photini Sinnis and Sean Prigge for MSP1, UIS4, and ACP antibodies, respectively. We also thank Dr. Puran Singh Sijwali (CCMB, Hyderabad) for the pET32a-PfAtg8 plasmid. We thank the CBRS facility of CDRI for providing the Maybridge library compounds. We also thank Dr. V.A. Nagaraj for the modified version of the GOMO-GFP-Luc plasmid originally obtained from Addgene (#60976) deposited by Dr. Olivier Silvie and Rohini Nandi for help with generating *P. berghei* Luc parasites. We acknowledge the THUNDER (BSC0102) and MOES (GAP0118) Intravital and Confocal microscopy facility of CSIR-CDRI. We thank Rima Ray Sarkar and Anil Kumar for their technical assistance with microscopy. We acknowledge the Advanced Research Computing (ARC) facility, University of Calgary, for the in silico experiments. AM and PNS were supported by the University Grants Commission and Department of Biotechnology, Government of India research fellowships. This manuscript is CDRI communication no. 145/2022/SM. The work was supported by the CSIR-CDRI in-house projects [IHP0011].

## Author Contributions

A.M. and S.M. conceived and led the study, performed data analysis, interpreted the results, and wrote the manuscript. A.M. performed most of the experiments. P.N.S. performed molecular docking. The *P. falciparum* drug assay was performed by S.A.H. All the authors have read and approved the final manuscript.

